# Capturing biodiversity complexities while accounting for imperfect detection: the application of occupancy-based diversity profiles

**DOI:** 10.1101/2020.09.07.285510

**Authors:** Jesse F. Abrams, Rahel Sollmann, Simon L. Mitchell, Matthew J. Struebig, Andreas Wilting

## Abstract

1. Measuring the multidimensional diversity properties of a community is of great importance for ecologists, conservationists and stakeholders. Diversity profiles, a plotted series of Hill numbers, simultaneously capture all the common diversity indices. However, diversity metrics require information on species abundance. They often rely on raw count data without accounting for imperfect and varying detection, although detectability can vary between species and study sites. Hierarchical occupancy models explicitly account for variation in detectability, and Hill numbers have been expanded to allow estimation based on occupancy probability. But agreement between occupancy and abundance-based diversity profiles has not been investigated.
2. Here, we fit community occupancy models to simulated animal communities to explore how well occupancy-based diversity profiles reflect true abundance-based diversity. Because we expect occupancy-based diversity to be overestimated, we further tested a novel occupancy thresholding approach to reduce potential biases in the estimated diversity profiles. Finally, we use empirical data from a megadiverse bird community to present how the framework can be extended to consider trait or phylogeny-based similarity when calculating diversity profiles.
3. The simulation study showed that occupancy-based diversity profiles produced among-community patterns in diversity similar to true abundance diversity profiles, although within-community diversity was overestimated with the exception of richness. While applying an occupancy threshold reduced this positive bias, this resulted in negative bias in species richness estimates and slightly reduced the ability to reproduce true differences among the simulated communities. Application of our approach to a large bird dataset revealed differential diversity patterns in communities of different habitat types. Accounting for phylogenetic and ecological similarities between species reduced diversity and its variability among habitats.
4. Our framework allows investigating the complexity of diversity for incidence data, while accounting for imperfect and varying detection probabilities, as well as species similarities. Visualizing results in the form of diversity profiles facilitates comparison of diversity between sites or across time. Therefore, our extension to the diversity profile framework will be a useful tool for studying and monitoring biodiversity.

## 1. Introduction

Biological diversity represents the variety of organisms or traits and plays a central role in ecological theory (Loreau et al., 2001; Tilman, Isbell, & Cowles, 2014). Mathematical functions known as diversity indices aim to summarize properties of communities that allow comparison among different regions, taxa, and trophic levels (Morris et al., 2014; Daly, Baetens, & De Baets, 2018). They are often used in conservation as indicators of the integrity or stability of ecosystems, and are, therefore, of fundamental importance for environmental monitoring and conservation (Morris et al., 2014). Diversity is, however, a generic term describing the complex multidimensional properties of a community. Any diversity index reduces these multidimensional properties to a single number (Morris et al., 2014), which is problematic (Daly, Baetens, & De Baets, 2018).

The most commonly used diversity indices are species richness, Shannon’s diversity H’ and Simpson’s diversity D; the latter two combine measures of richness and abundance, whereas species richness solely presents the number of species. It is not uncommon that diversity increases according to one index, but decreases according to another (Patil, 2014), demonstrating the difficulties in quantifying biodiversity in a single number (Purvis & Hector, 2000; Daly, Baetens & De Baets, 2018).

To address this shortcoming, several researchers have suggested using parametric families of diversity indices (Hill, 1973; Patil & Taillie, 1982; Gattone & Battista, 2009; Leinster & Cobbold, 2012). Jost (2006) proposed the use of Hill numbers (Hill, 1973) that incorporate relative abundance and species richness to show the number of equally abundant species necessary to produce the observed value of diversity. Individual Hill numbers differ by the parameter *q*, which quantifies how much the measure discounts rare species when calculating diversity (the higher *q*, the less these rare species contribute to diversity). Plotting the effective number of species as a function of *q* allows us to view diversity from multiple vantage points (Hill, 1973; Leinster & Cobbold, 2012) and the resulting curves have become known as diversity profiles. These curves display different properties of diversity and often drop sharply between *q*=0 and *q*=1 and level off soon after *q*=2, indicating that many communities are dominated by few highly abundant species (Preston, 1948).

To calculate diversity indices (except for species richness), as well as diversity profiles, information about the relative abundances of species, or evenness, is required (Leinster & Cobbold, 2012). Obtaining information on species abundance can, however, be challenging. Raw count data are typically fraught with detection bias (Nichols et al., 1998; MacKenzie & Kendall, 2002; Sollmann et al., 2013), and inference about diversity from indices based on count-based relative abundance estimates that do not account for imperfect and varying detectability may therefore be biased. Estimating abundance of all species in a community while accounting for varying detection, for example using capture-recapture methods (Royle & Dorazio, 2008), is extremely difficult as different organisms require different sampling methods to obtain sufficient data for reliable abundance estimation. Thus, community studies often resort to the collection of much cheaper and easier to obtain detection/non-detection data. Even with incidence data, however, we must consider that species may be detected imperfectly (MacKenzie et al., 2002; MacKenzie et al., 2006).

Occupancy modelling provides a framework to handle the problem of imperfect and varying detection, producing unbiased estimates of species occurrence (MacKenzie et al., 2002, 2006). The development of hierarchical multi-species occupancy models (Dorazio & Royle, 2005; Dorazio et al., 2006) has enabled estimation of richness at the level of the study area and survey location (Sollmann et al., 2017), and to model variation in richness across areas as a function of covariates (Sutherland et al., 2016). However, it has been shown that multi-species occupancy models overestimate true species richness (Zipkin et al., 2012), and only a few applications for other diversity indices exist (Guilleral□Arroita, Kéry, & Lahoz□Monfort, 2019). Consequently, accounting for imperfect detection has often been neglected in calculating diversity metrics in the past.

Only recently, Chao et al. (2014) described a method to calculate Hill numbers from incidence data, and Broms, Hooten, & Fitzpatrick (2014) followed with a formulation to calculate occupancy-based Hill numbers. Here, we extend this framework to facilitate the calculation, visualization, and thus, interpretation of occupancy-based diversity profiles. We first explore how well occupancy-based diversity profiles reflect true abundance-based diversity and their ability to compare communities across landscapes with varying levels of habitat disturbance using simulated data. Because occupancy is a coarser measure than abundance, containing less information about how relatively abundant or rare a given species is in its community, we expect occupancy-based profiles to overestimate diversity for *q*>0 (i.e., suggest a more even community); but because this affects all communities, we expect occupancy-based profiles to be able to correctly order areas by their diversity. We then used an empirical dataset of diverse bird communities collected in Sabah, Malaysian Borneo, to demonstrate how the framework can be extended to a trait-based diversity analysis of occupancy-data by incorporating measures of similarity as proposed by Leinster and Cobbold (2012).

## 2. Methods

Our study was divided in into a simulation study and the application of the occupancy-based diversity profiles to an empirical dataset. The simulation study followed seven steps: (1) simulation of a forest degradation gradient, (2) simulation of animal communities, and detection data of those communities, (3) community occupancy analysis of simulated detection data, (4) utilization of community occupancy model output to construct occupancy-based diversity profiles, (5) application of thresholding to occupancy-based profiles, and (6) evaluation of the performance of the occupancy-based diversity profiles by comparing them to true community abundance diversity profiles. In the second part of our study we applied the community occupancy diversity profiles to an empirical bird dataset from Malaysian Borneo and tested how these profiles could account for phylogenic or ecological trait similarities.

### 2.1 Forest degradation and community simulation

We simulated five virtual 10 × 10 km forest landscapes, with 200 × 200 m grid cells (50 × 50 cells). We simulated a habitat covariate representing “habitat disturbance” (where 0 represents undisturbed forest and 5 represents complete deforestation) for each landscape (Fig. 1A) by drawing random samples from a multivariate normal distribution. To increase realism by simulating nonrandom habitat, we explicitly included spatial autocorrelation in the simulation of the habitat covariate by using the distance matrix as our variance-covariance matrix and a decay function with a decay constant *φ* to specify how the relationship to other cells changes with distance (Suppl. Fig. S1). The five landscapes constitute a habitat degradation gradient representative of three different logging regimes and two “patchy” landscapes that simulate activities such as compartmental logging: (1) no disturbance, (2) patchy low disturbance, (3) low disturbance across the entire area, (4) patchy high disturbance, and (5) high disturbance across the entire area.

**Figure 1.**
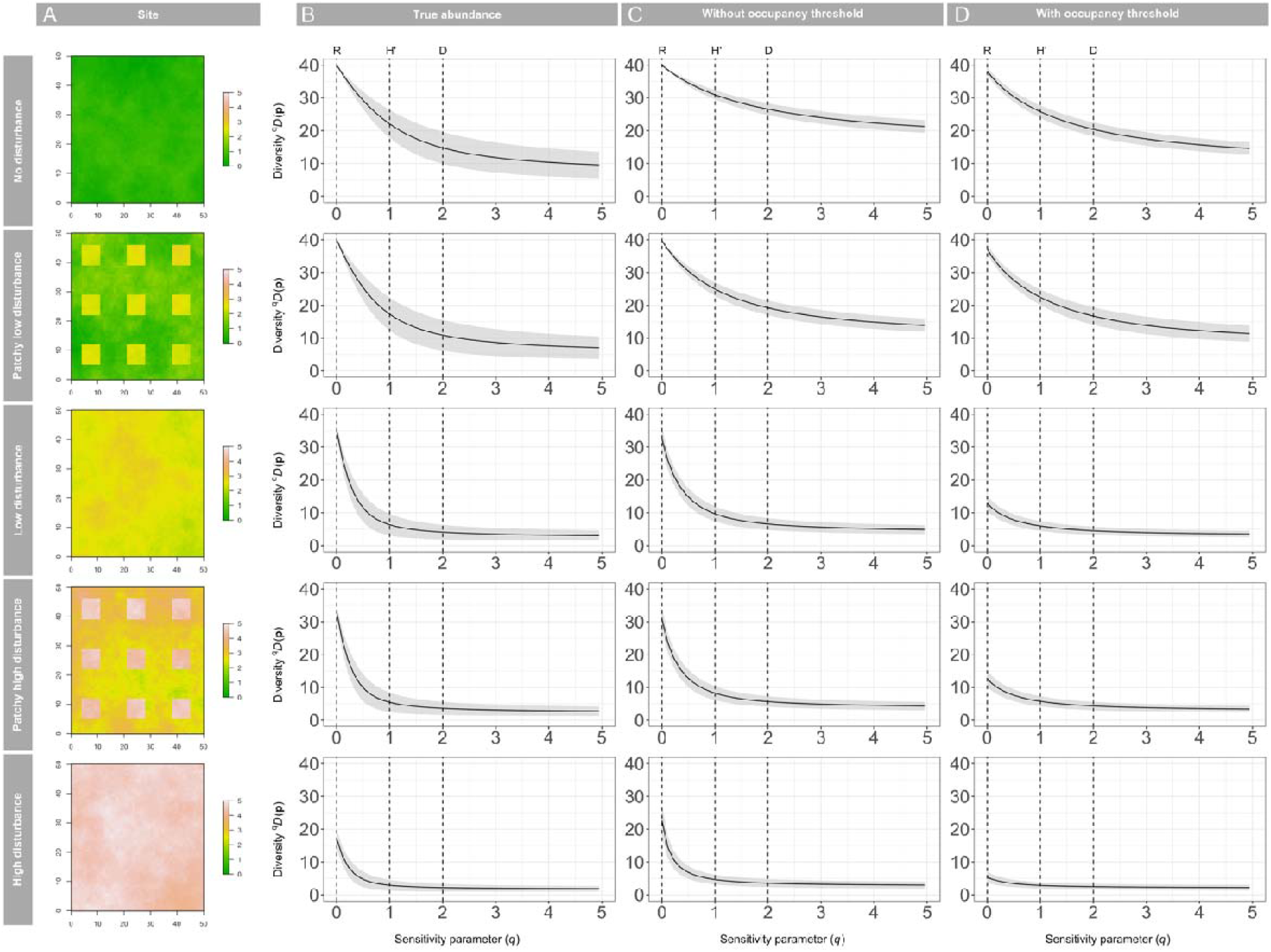
Results from the simulation study. **(A)** Simulated “forest quality” habitat covariate for five virtual landscapes. Average diversity profiles (solid line)and standard deviation (grey shading) for simulated data generated using **(B)** the simulated true abundance, and the community occupancy predictions for the entire landscape **(C)** without thresholding and **(D)** with thresholding using the max_SSS_ method for the entire study landscape.

We then simulated 100 communities for each landscape. We simulated the cell-level abundance of forty virtual species, *i*, at grid cell *j* (*j* = 1, 2,…,*n*) to generate “true” abundance-based diversity profiles. We set the average response of species to habitat disturbance as negative (µ_1_ = −2) since most forest-adapted species will respond to logging negatively (Suppl. Fig. S2). We allowed this to vary, generating a community of species with mostly negative responses, but with few species that responded positively to habitat disturbance. Average expected abundance per species per grid cell in undisturbed forests was 0.37. This resulted in five communities with species richness ranging from 17.2 (±2.8) for the most disturbed site to 40 (±0) for the undisturbed site. See Supplementary materials for full simulation details.

For each simulated community we then generated detection/non-detection data by simulating systematic repeated sampling of the community. For each landscape, we picked 100 sampling points in a grid spaced 2 km apart. At each point, we then simulated the observation process with imperfect detection. We repeated the observation process for 10 occasions, *k*. Species-specific detection on the logit scale, *α*0, was drawn from a normal distribution with *µ* _*α*_ = 0 and *σ* _*α*_ = 1.

Finally, we reduced the resulting observed counts to detection/non-detection data. We chose to represent the detection process in two steps (generating counts, then reducing these to binary detection/non-detection data) to reflect how data from typical non-invasive survey methods, such as bird point counts or camera-trapping, are prepared for occupancy modeling. We constructed and saved abundance diversity profiles for later comparison to the occupancy-based diversity profiles. All calculations were carried out in R version 3.6.0 (R Core Team, 2019).

### 2.2 Community occupancy model

We adopted the hierarchical formulation of occupancy models by Royle & Dorazio (2008) extended to a community occupancy model (Dorazio & Royle, 2005; Dorazio et al., 2006). To analyze our simulated data, following common practice in analyzing field data, we modelled occupancy probability as having species-specific random intercepts, *β*0_*is*_, with landscape specific (indicated by *s* indexing) hyperparameters (*μ*_*β*0,*s*_, *σ*_*β*0,*s*_), to allow for species-specific effects on occupancy of the simulated habitat “disturbance” covariate (*β*1_*i*_). different baseline occupancy in the reserves and among species. We further modelled Detection probability included a species-specific random intercept with landscape specific hyperparameters, to allow for differences in baseline detection among reserves. In our case, the different landscapes had different abundances of animals, which leads to differences in species-level detection (Royle & Nichols, 2003). The formal model description can be found in the supporting information. We implemented the model in a Bayesian framework using JAGS (Plummer, 2003) accessed via the R packages *rjags* (Plummer, 2019). We ran three parallel Markov chains with 250,000 iterations, of which we discarded 50,000 as burn-in, and we thinned the remaining iterations by 20 to make the output more manageable. We assessed chain convergence using the Gelman-Rubin statistic (Gelman et al., 2004). Values under 1.1 indicate convergence, and all parameters in our models had a Gelman-Rubin statistic <1.1. We tested whether the model adequately fit the data by calculating a Bayesian p-value (Gelman et al., 1996).

### 2.3 Diversity profiles

In order to investigate the performance of estimating diversity profiles with occupancy-based information, we first constructed diversity profiles for the simulated true abundance across each landscape, following Leinster & Cobbold (2012). Diversity profile values (^*q*^*D*^*Z*^) for abundance data can be calculated according to Leinster & Cobbold (2012) as:

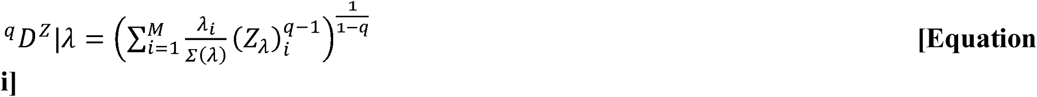

where

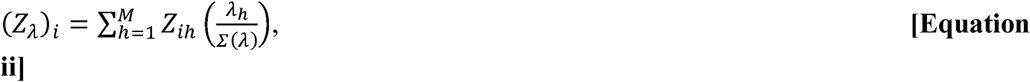

*M* is the number of species in the assemblage, and the *i*th species has relative abundance 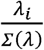. The parameter *q* determines the sensitivity of the measure to the relative abundances of species. This allows us to calculate diversity along a continuum of values of *q*. At *q*=0, ^*q*^*D*^*Z*^ equals species richness where all species are considered equally. As *q* becomes larger, more weight is placed on common species thereby incorporating evenness into the diversity measure and resulting in a lower value of ^*q*^*D*^*Z*^ for more uneven assemblages than for more even assemblages. The diversity profile framework from Leinster & Cobbold (2012) allows similarity matrix **Z** which represents the similarity between the *i*th and *h*th species. Values of for the consideration of similarity between species through the inclusion of an *M* x *M* 0 in **Z** indicate total dissimilarity, whereas values of 1 indicating identical species. This matrix can be used to adjust the profiles by incorporating any measure of similarity (such as phylogenetic or trait) between different species or taxonomic groups. In our simulation study, we use a naive similarity matrix (an identity matrix with all cells on the diagonal equal to 1 and all other values = 0). In the empirical dataset (see below) we adjusted the profiles using a diet, taxonomic, and a phylogenetic similarity matrix. We refer to *q*=0 as “richness” (R), which, depending on the nature of *Z*_*ih*_, can represent species richness, or trait richness.

To use occupancy probabilities instead of abundances to construct diversity profiles we altered the diversity profile method of Leinster & Cobbold (2012) as follows:

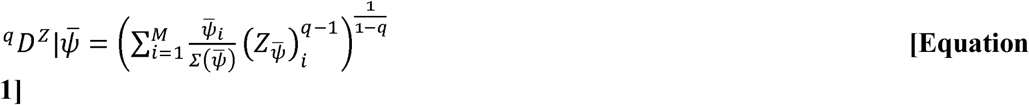

where

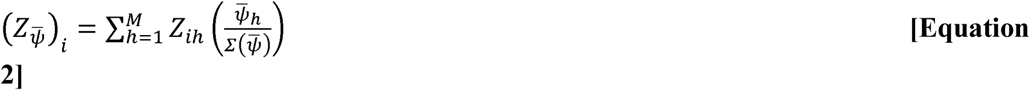

where *M* is again the number of species in the assemblage, 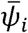 is the average occupancy probability of species *i*, the *i*th species has relative occupancy 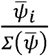, and **Z** is the similarity matrix.

We used the parameter estimates from the occupancy model fit to our simulated data to predict species distributions across the five simulated landscapes (i.e., not just the sampling points) and then constructed diversity profiles using the mean landscape-wide occupancy probability for each species using Equations 1 and 2 for all posterior samples of the community occupancy model. This effectively creates posterior distributions for the diversity profiles themselves and allowed us to determine their standard deviations and 95% credible intervals. We then compared estimates of species richness (R), Shannon’s diversity H’ and Simpson’s diversity D, from the occupancy-based profiles against indices based on the true abundance profiles. Specifically, for each landscape, we present the average relative bias (occupancy-based index minus true abundance index divided by true index) across all 100 communities and coverage, i.e., the proportion of communities for which the 95% BCI of the occupancy-based index estimate included the true-abundance based index.

To evaluate how well occupancy-based diversity profiles were able to order landscapes by site diversity rank, we compared them to the diversity ranking in the true abundance-based profiles. Since not all communities were different in the true abundance based profiles (e.g. no disturbance and patchy low disturbance landscapes both had an average species richness of 40 (Table 1) and could, therefore, not be distinguished in the true-abundance based profiles), we first determined how many sites we could reliably distinguish. To do so we used the results of the true abundance-based simulations and checked which landscapes could be distinguished in 95% of the 100 simulations for R, H’, and D. We were able to distinguish 3, 4, and 3 of the 5 sites in the true abundance profiles for R, H’, and D, respectively (see Suppl. Table S1). We then calculated the proportion of communities for which occupancy-based diversity profiles resulted in the same rank order of these distinguishable landscapes as true abundance-based profiles.

**Table 1.**
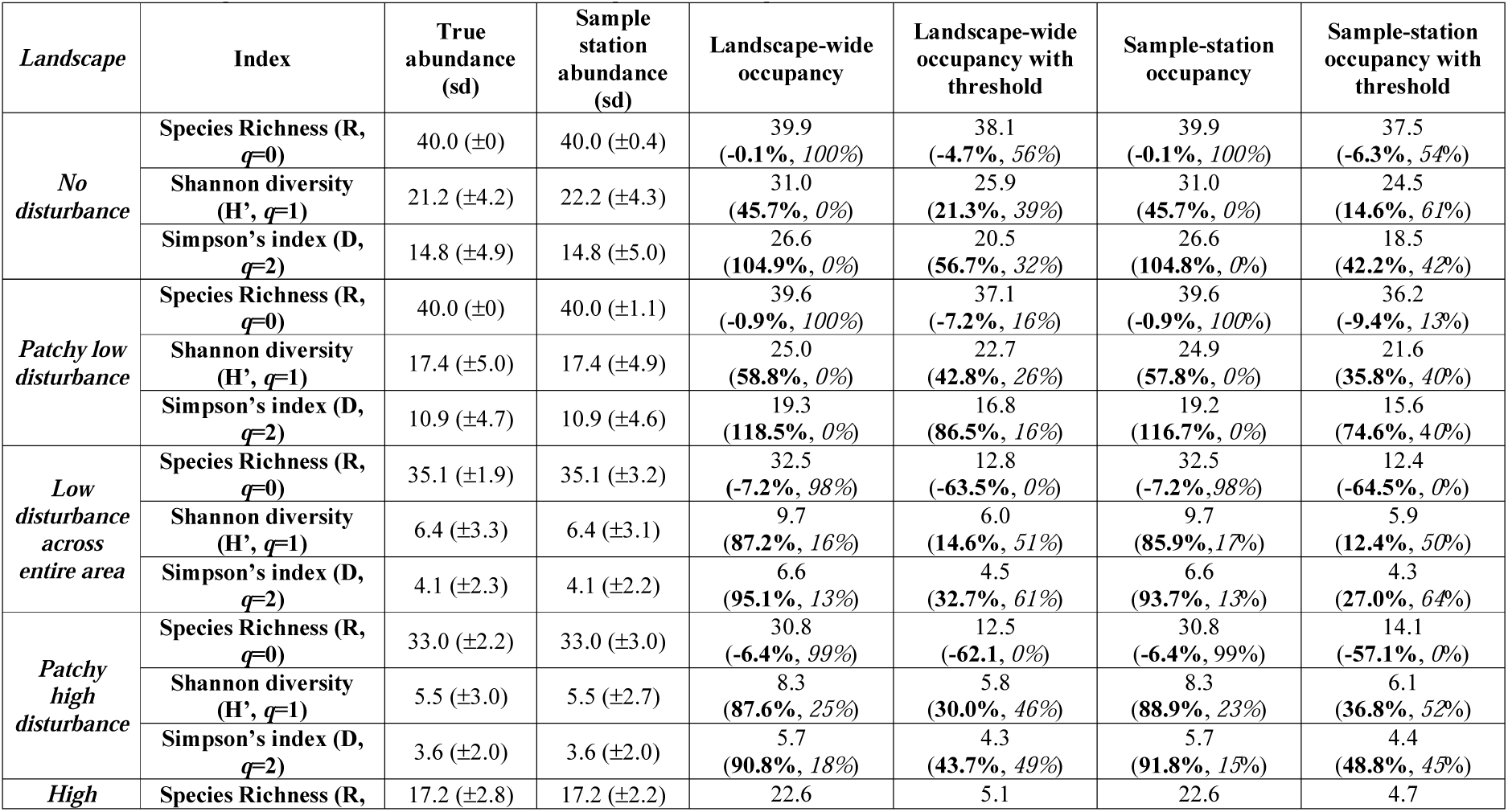
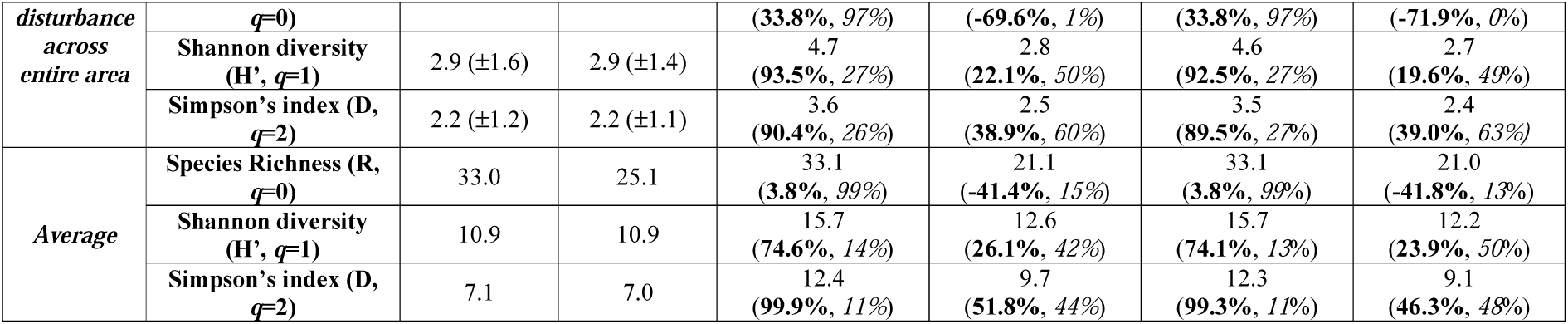
Summary of occupancy-based diversity index estimates for 5 simulated landscapes. Parentheses indicate the relative bias (bold) and 95% BCI coverage (italics) averaged over the 100 simulations when compared to the respective true abundance-based index. For the true and sample station abundance-based indices, parentheses represent the standard deviation over the 100 simulations.

In some practical applications of this method, it may not be possible to predict occupancy across the entire landscape (missing covariates for unsurveyed cells). Therefore, we also calculated diversity profiles for both true abundance and occupancy data for just the sampling points and compared these to the landscape scale profiles.

### 2.4 Occupancy Threshold

To explore methods to account for the expected overestimation of diversity profiles in community occupancy models, we tested the use of an occupancy threshold. When using presence/absence data, the identities of both presence and absence data are (assumed to be) known (Liu et al., 2016). However, with detection / non-detection data we have no information about “true absences”, which presents challenges for threshold selection. Liu et al. (2016) identified the *max*_*SSS*_ method, which maximizes the sum of sensitivity and specificity, as the most suitable objective approach for determining thresholds with incidence data.

The *max*_*SSS*_ method described by Liu et al. (2016) requires the use of “pseudo-absences”, which are randomly picked from the sampling stations with no detections. Here, we use the estimates from the occupancy model to draw “pseudo-absences” randomly for stations without detections, but weighed by the probability of a station being unoccupied, 1 − *ψ*.

We calculated the *max*_*SSS*_ (Suppl. Fig. S3) threshold for each species for each landscape based on the mean occupancy estimate using the *optimal.thresholds* function from the R package *PresenceAbsence* (Freeman & Moisen, 2008). We set occupancy probabilities for stations with estimates below the occupancy threshold to zero. We then averaged the threshold-adjusted occupancy for each species across landscapes for each model iteration and generated new diversity profiles using the adjusted dataset. We compared threshold occupancy-based profiles against true-abundance based profiles as described for non-threshold profiles.

### 2.5 Case study

We sampled bird communities at 307 point-count localities in and around the Stability of Altered Forest Ecosystems (SAFE) project (117.5°N, 4.6°E) in Sabah, Malaysian Borneo (Mitchell et al., 2018). Thirty-eight localities were in continuous logged forest (CF) of the Ulu Segama Forest Reserve, with an additional 156 in the neighboring SAFE landscape, in forest that had been logged several times and recently salvage logged. A further 113 localities were sampled alongside rivers in oil palm plantations, including 88 with riparian forest remnants (RR) on each riverbank and 15 with no natural vegetation (OPR). Localities are classified by habitat into four categories: non-riparian continuous forest (CF), riparian forest (RF), riparian remnant (RR), and oil palm river (OPR). Forest quality, based on aboveground carbon density measured via LiDAR, also varied substantially across the landscape.

We observed a total of 169 bird species. Two species, *Leptocoma brasiliana* and *Zanclostomus javanicus*, were excluded because there is no phylogenetic information available, which is necessary for the trait-based analysis. Further, three species of swift (*Aerodramus maximus, A. salangana* and *A. fuciphagus*) could not be reliably separated and are considered as *Aerodramus spp*.

To analyze the case study data, we used a similar community occupancy model structure as used for the simulation study. Following Mitchell et al. (2018), we modeled occupancy using above-ground carbon density, forest cover and riparian remnant width as predictors, with species-specific random intercepts with habitat-specific hyperparameters. Covariates were derived using remotely sensed data and calculated following Mitchell et al. (2018). Detection probability included a species-specific random intercept with habitat specific hyperparameters and accounted for the effect of time and date of a survey on the probability of detection (e.g., Ellis & Taylor, 2018).

We separated species communities according to the four habitat-types described above for diversity profile construction with and without occupancy thresholds. We did not do landscape scale predictions for the case study, but instead used the occupancy probabilities for each sampling station to construct the diversity profiles. Additionally, we constructed similarity matrices (see Supplementary Material) according to diet, taxonomy, and phylogeny to demonstrate how similarity can be incorporated into our occupancy-based diversity profile framework.

## 3. Results

### 3.1 Simulation results

We present the average and standard deviation for the true abundance and occupancy-based profiles generated for the 100 simulated communities in Figure 1. The predicted occupancy-based diversity profiles (Fig. 1C without thresholding and 1D with the threshold for the application of this method to a single site see Suppl. Fig. S4) show similar trends in diversity as the diversity profiles based on true abundance (Fig. 1B). The occupancy-based diversity profiles, however, often overestimated diversity, particularly for *q* > 0.5. Whereas species richness estimates without thresholding corresponded very well to true species richness (−0.1 to −7.2% average bias) for all but the “high disturbance across entire area” landscape (average bias of 38%), H’ and D were consistently overestimated (Table 1 and Fig. 1B). Applying the threshold reduced this overestimation, but at the same time resulted in an underestimate of species richness, particularly in more disturbed landscapes (Table 1).

For all five landscapes, at values of *q* > 0 there was a positive bias in the non-threshold occupancy-based profiles/diversity indices when compared to the true abundance profiles/indices (Table 1, Fig. 1). This positive bias was stronger for *q* > 1 than for *q* < 1. While the Shannon diversity H’ was overestimated by between 45.7% (no disturbance landscape) and 93.5% (high disturbance across the entire landscape), Simpson’s index was overestimated by between 90.4% (high disturbance across the entire landscape) and 118.5% (patchy low disturbance). There was a tendency for bias in H’ to increase with increasing disturbance, whereas bias in D tended to decrease in more disturbed areas. Patterns in coverage mirrored patterns in bias, with nominal (>95%) coverage of richness, but poor coverage (between 0 and 27%) of the other two indices (Table 1).

The application of an occupancy threshold (Fig. 1C) resulted in negative bias in estimates of species richness, especially for the three more disturbed landscapes in which species richness was underestimated by between 63.5% and 69.6% (Table 1). In these three more disturbed sites species had lower occupancies; communities were, therefore, more vulnerable to the exclusion of some species after application of the threshold. For the two less disturbed landscapes the threshold-based estimates of richness were comparable to the true abundance estimates (Table 1 and Fig. 1C). The threshold improved the agreement between the occupancy and abundance-based profiles for values of *q* > 0. Bias in estimates of H’ after threshold application ranged from 21.3 to 42.8%, and for D from 32.7 to 86.5%. The occupancy threshold reduced coverage for richness to between 56% in the undisturbed landscape and 0% in the most disturbed landscape. The threshold increased coverage for H’ and D to between 16 and 60%, with a tendency for coverage to increase with increasing disturbance.

For both abundance and occupancy-based profiles, we did not find great discrepancies between the landscape-wide profiles and the sampling station-based profiles, even though the sampling stations only covered about 4% of the study area (Table 1, Suppl. Fig. S5).

For the comparison across landscapes the occupancy-based diversity profiles generally maintained the same rank order of diversity amongst the landscapes (Fig 2A and 2B, for an example of results from a single site see Suppl. Fig. S6) as the true abundance-based diversity profiles. The three communities that showed significantly different species richness based on true abundance data were ordered the same by occupancy-based richness estimates in 97% of the simulated communities (Fig. 2C and 2D, Table 2). Similarly, the four and three landscapes that could be significantly distinguished for H’ and D were ordered the same as occupancy-based profiles in 96% and 94% of all simulations, respectively. The occupancy threshold slightly increased the ability to correctly rank simulated landscapes by species richness (to 100%), but slightly reduced it for H’ and D (Fig 2E & 2F, Table 2).

**Table 2:** Percentage of the 100 simulated communities within each of five landscapes where occupancy-based diversity profiles were ordered the same as the true abundance-based diversity profiles. The maximum number of landscapes that could be reliably separated in the true abundance profiles was 3, 4, and 3 of the 5 landscapes for R, D, and H’, respectively (see Suppl. Table S1).

| Index | Landscape-wide occupancy | Landscape-wide occupancy<br>with threshold | Sample-station occupancy | Sample-station occupancy<br>with threshold |
| --- | --- | --- | --- | --- |
| Species Richness (R,<br>*q=0*) | 97% | 100% | 97% | 100% |
| Shannon diversity<br>(H', *q=1*) | 96% | 94% | 96% | 92% |
| Simpson's index (D,<br>*q=2*) | 94% | 90% | 94% | 82% |

**Figure 2.**
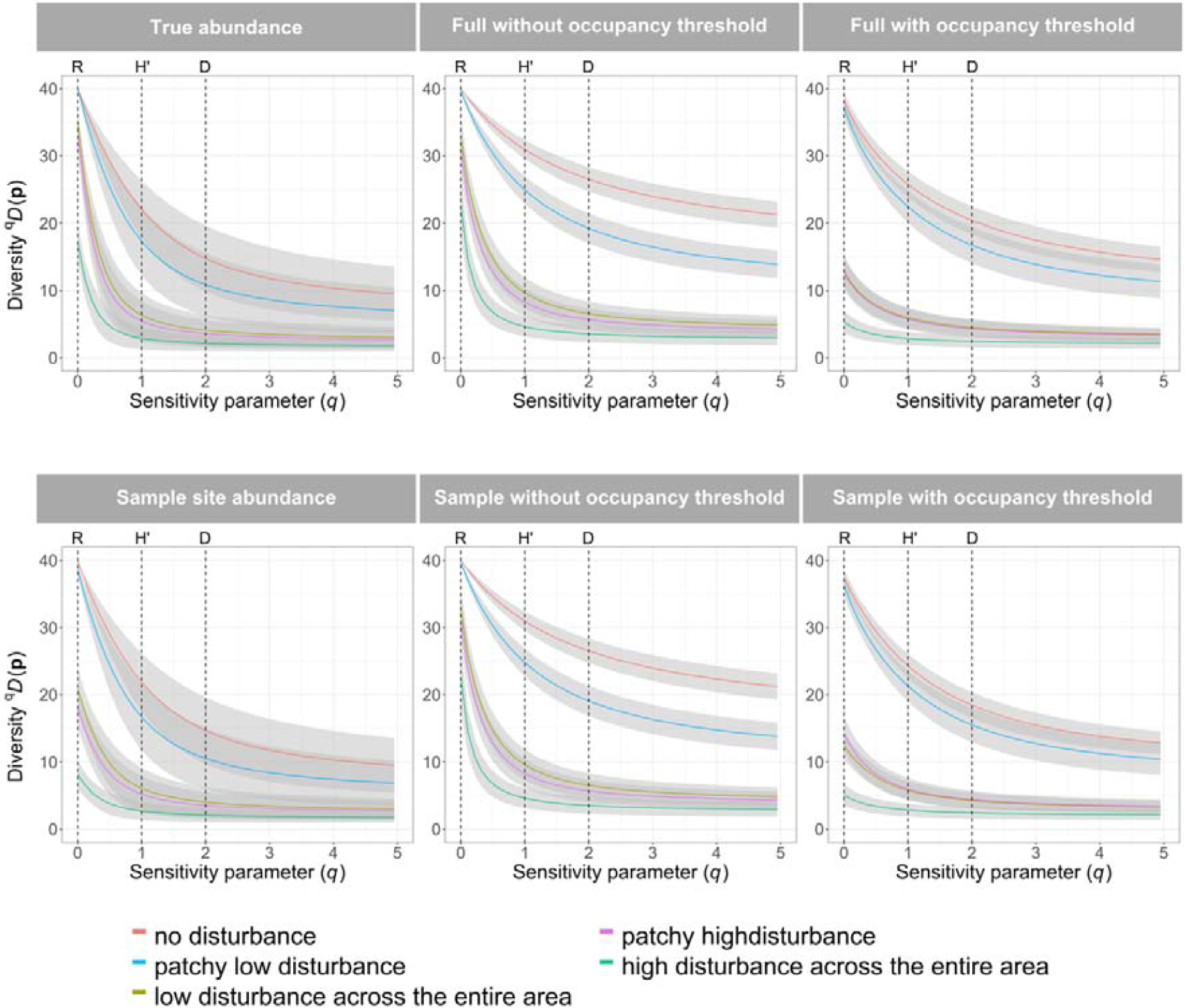
Comparison among landscapes of diversity profiles generated using **(A)** the true abundance across the whole landscape, **(B)** the true abundance at the 100 sample stations in each landscape, **(C)** occupancy based predictions across the whole landscape without thresholding, **(D)** occupancy based predictions at the 100 sample stations in each landscape without thresholding, **(E)** occupancy based predictions across the whole landscape with thresholding, **(F)** occupancy based predictions at the 100 sample stations in each landscape with thresholding. All results are shown as averages (solid line) of the 100 simulations with the uncertainty shown as standard deviations (grey shading).

### 3.2 Borneo bird community results

We detected 143, 118, 121, and 30 species in continuous forest, riparian forest, riparian remnant, and oil palm river, respectively. In the diversity profiles, species richness was highest in continuous forest, followed by (in decreasing order) riparian remnant, riparian forest, and oil palm river. At *q* < 1, continuous forest was the most diverse habitat type, while at *q* > 1.5 riparian forest was the most diverse (Fig. 3A). Although the 95% BCIs of the profiles for these two more diverse habitats overlapped, this indicates that the continuous forest community is the richest in species but that this community is less even than the riparian forest community. Although species richness in the riparian remnant was similar to continuous forests and riparian forest, the diversity decline of the profile was much greater, so that riparian remnants were significantly (non-overlapping 95% BCIs) distinct from continuous forest and riparian forest at *q* > 0.5. Oil palm river and riparian remnant were always the habitats with the lowest and second lowest biodiversity, respectively, and at *q* > 2 the 95% BCIs of these two profiles largely overlapped.

**Figure 3.**
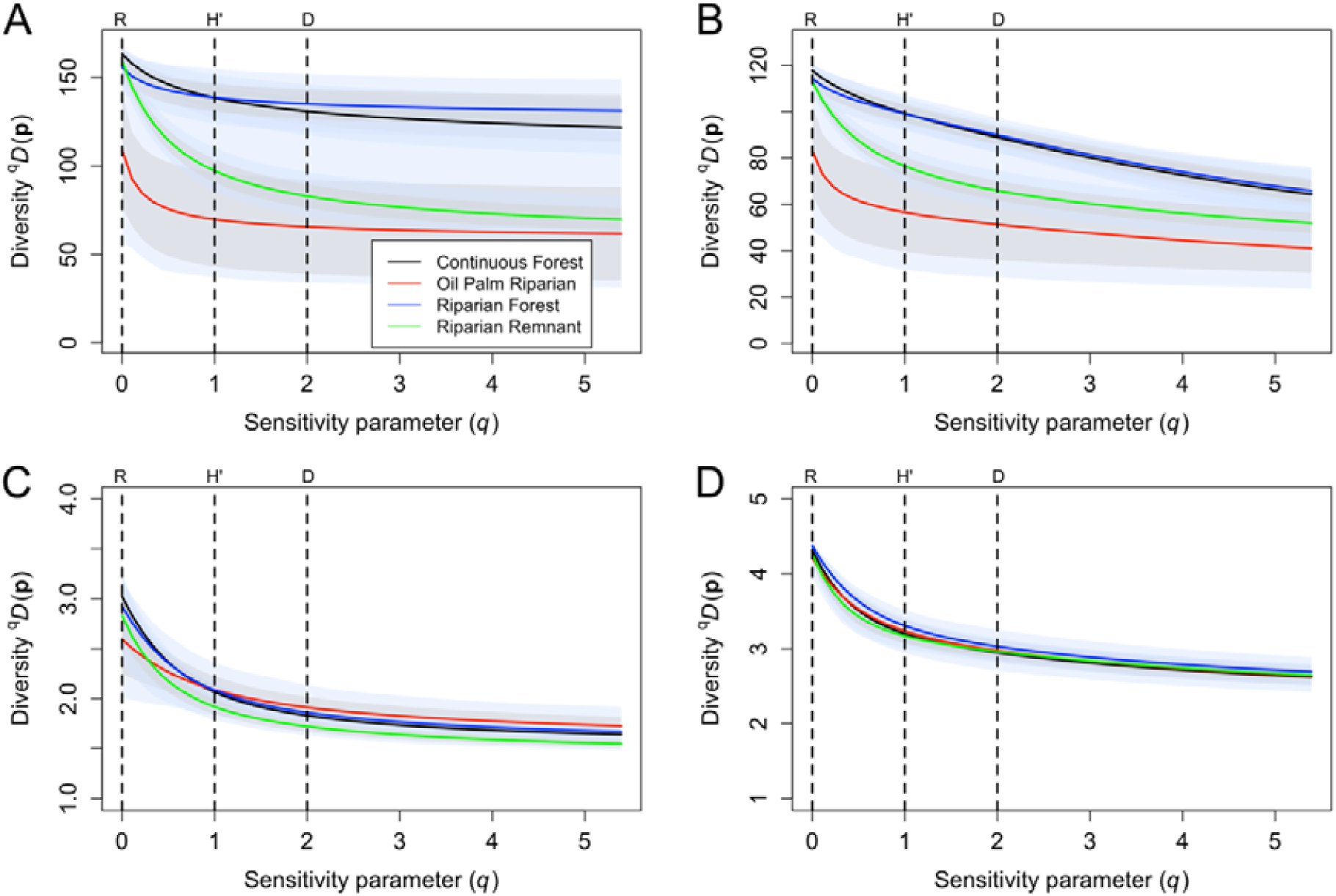
Diversity profiles for a bird community from Malaysian Borneo in four habitat types calculated using the occupancy predictions without thresholding. **(A)** naive similarity matrix, **(B)** taxonomic similarity, **(C)** diet similarity, **(D)** phylogenetic similarity. The standard deviations (grey shading) and 95% credible intervals (blue shading) are included to quantify uncertainty.

When we incorporated similarity (diet, taxonomic, and phylogenetic), the overall community diversity was reduced for all habitat types. In line with species richness, the taxonomic richness (Fig. 3C) was similar for the riparian remnant, continuous forest and riparian forest. Continuous forest and riparian forest showed very similar overlapping profiles. The shape of the riparian remnant and oil palm river taxonomic profiles were very similar to the original profiles but overlap in the 95% BCIs was even greater. When we considered dietary (Fig. 3B) and phylogenetic (Fig. 3D) similarity, we also saw a large reduction in the diversity of the communities. In both cases, all habitats showed very similar diversity profiles with widely overlapping 95% BCIs.

With thresholding (see Suppl. Fig. S7), the estimated species richness was lower for all habitat types. Profiles with the threshold showed similar patterns, but differences among habitats, particularly for the more depleted communities in the riparian remnant and the oil palm river, were less distinct.

## 4. Discussion and Conclusion

Diversity profiles allow researchers to characterize and compare communities while considering the contributions of abundant and rare species, thus acknowledging the multidimensional nature of diversity (Morris et al., 2014). We aimed to provide a reliable inference framework for estimating biodiversity via diversity profiles based on output from occupancy models, which take into account imperfect and varying species detection. As expected, we found that occupancy-based diversity profiles generally mirrored true among-community patterns well, albeit with some shortcomings that are primarily related to overestimations of non-richness diversity measures.

Using occupancy model estimates of average occupancy probability to construct diversity profiles, with or without thresholding, generally maintained the same inter-landscape diversity pattern as observed in the true abundance diversity profiles (Fig. 2). Further, the two simulated landscapes that had very similar abundance-based profiles (landscape-wide low disturbance and local high disturbance) also showed very similar occupancy-based profiles. The general agreement of the occupancy and true abundance profiles suggests that detection/non-detection surveys may be sufficient to compare the multidimensional properties of diversity between landscapes. Similarly, the repeated collection of detection/non-detection data from one landscape will likely allow comparison of diversity through time, an important aspect of biodiversity monitoring.

Within a landscape, occupancy-based profiles generally overestimated diversity for *q*>0. This is expected as occupancy probability is bound between 0 and 1, whereas abundance has no upper bound; consequently, occupancy-based diversity should suggest more even communities (i.e, be higher) than abundance-based diversity. Further, Broms, Hooten, & Fitzpatrick (2014) also found positive bias in when comparing their occupancy-based to true incidence-based Hill numbers and attributed that to positively biased estimates of occupancy in species with low detection rates. This bias is caused by the structure of the community occupancy framework, as rare species borrow information from common species (Guillera□Arroita, Kéry, & Lahoz□Monfort, 2019), and may also contribute to the positive bias we observed. Similar to Broms, Hooten, & Fitzpatrick (2014), we saw no or low negative bias in estimates of species richness (*q*=0), with exception of the most disturbed landscape, which had 33.8% positive bias in richness estimates, on average. This most likely relates to the sparsity of many of the species in this landscape, translating into low detection probabilities and thus, sparse data, and leading to small sample bias.

Acknowledging the differences between abundance and occupancy, we do not suggest that occupancy probability can simply be used as an index or surrogate of abundance. Our results do, however, indicate that under conditions representative of the methods commonly used in wildlife research, occupancy-based diversity measures and profiles can reflect patterns in diversity despite the loss of information entailed in using occupancy rather than abundance data. This is a promising finding for biodiversity research and monitoring, as community-wide species detection/non-detection data are generally much easier and cheaper to obtain than data for abundance estimation (Joseph et al., 2006; Kéry & Schmidt, 2008). In addition to traditional methods used to detect wildlife, such as points counts and visual transects, a growing number of technologies are available for detecting and identifying biodiversity, such as automated acoustic recorders (Bush et al., 2017) or eDNA and metabarcoding (Bush et al., 2017). These new technologies present powerful methods to collect community level incidence data that could be combined with occupancy-based diversity profiles.

We attempted to overcome the overestimation of diversity from community occupancy models by implementing an occupancy threshold. Liu et al. (2016) suggest the use of randomly selected points as pseudo-absences for threshold determination with incidence data. Here, we used the output of the occupancy model to generate pseudo-absences based on estimated occupancy probability. This approach likely leads to more realistic pseudo-absences than completely random generation as the use of modeled occupancy probabilities allows for a more informed selection. Incorporating a threshold into occupancy-based diversity profile calculation had mixed results. The threshold reduced the overestimation of diversity at *q* > 0.5 (Table 1). At the same time, however, thresholding often resulted in underestimated species richness, particularly for more disturbed landscapes (Table 1) for *q* > 0. Interestingly, the use of a threshold did not improve, but rather slightly reduced our ability to correctly replicate true underlying differences in diversity among landscapes (Table 2). In the empirical dataset, we saw a similar reduction in the distinctiveness of diversity profiles for the riparian remnants and the oil palm rivers, whose 95% BCIs largely overlapped for *q* > 1 when a threshold was applied. The application of the threshold therefore depends on the research question and the diversity within the community. If the goal is the comparison of profiles between sites or of the same site across time, which we expect is the main objective of most biodiversity research and monitoring projects, we recommend to not use a threshold. In this case, researchers should be aware that species richness (when data are sparse for many species) as well as the overall diversity profile at *q* > 0.5 (particularly in undisturbed more diverse landscapes) may be overestimated. We acknowledge that we only explored one threshold and that effects may be different for other threshold methods.

Occupancy-based diversity profiles derived from landscape scale predictions were comparable with occupancy-based profiles using only the sampling stations, which covered 4% of our simulated landscapes. This indicates that reliable profiles can be generated in situations where landscape-wide predictions are not possible due to missing covariate information at unsampled locations. However, this only holds when sampling is representative of the entire landscape. The risk of non-representative sampling is higher in more heterogeneous and more logistically challenging landscapes or if the number of stations is much lower (the simulated 100 stations is a high number for many field projects).

The diversity profile framework presented here also allows for the incorporation of trait similarities between species by defining a similarity matrix. Incorporating species trait similarities can be an additional way to display diversity in a community as it puts a greater emphasis on more dissimilar species (Leinster and Cobbold, 2012). From an ecological perspective, accounting for such similarities reduces the functional redundancies in the community, for example, species having the same dietary niche could functionally replace each other (Rosenfeld, 2002; Olden et al., 2004). Phylogenetically and functionally diverse communities are known to better maintain ecosystem stability (Cadotte, Carscadden & Mirotchnick, 2011; Cadotte, Dinnage & Tilman, 2012). Therefore, considering these additional dimensions of diversity provides a more complete picture of a community (Rodrigues & Gaston, 2002) and may improve predictions of ecosystem function and resilience.

Our empirical data showed that considering dietary, taxonomic or phylogenetic similarities among bird species led to very similar diversity profiles for all habitat types. In the case of this bird dataset, all taxonomic, phylogenetic and dietary groups were present in all habitats. As a result, even at considerably lower species diversity, disturbed habitats such as oil palm plantations maintained dietary and phylogenetic diversity of birds essentially identical to that of continuous forests. This is surprising, given that previous studies have found that dietary traits and taxonomy (among other characteristics) can affect response to habitat alterations and extinction risk in birds (e.g., Russell et al. 1998; Boyer 2010; Frishkoff et al. 2014). Despite this apparent maintenance of phylogenetic and functional diversity, the loss of overall species diversity in more disturbed habitats suggest a loss in redundancy, another measure that has been associated with ecosystem stability (Naeem, 1998).

The diversity profiles of the bird communities reinforced the findings by Mitchel at el. (2018) that riparian remnants supported similar diversity value to continuous logged forest habitats (both riparian and non-riparian). However, when evenness of the community is given more weight (i.e. when *q* > 1), riparian remnants have reduced diversity compared to logged forests. This suggests that much of the diversity in the habitat remnants (such as when measured via species richness directly), manifests from a number of species occurring rarely. If a greater proportion of the community in remnant occurs only rarely, this suggests such remnants may not sustain certain species in the long-term (i.e., we may be observing an extinction debt) and effectively act as population sinks from continuous forest habitats, a finding which is not apparent from assessing only species richness and community integrity as undertaken by Mitchell et al. (2018).

In practice, information on species occurrence is often used to help develop management decisions and conservation strategies (Guisan et al., 2013). For many species of conservation concern, the detection/non-detection surveys underlying estimates of occurrence are the main source of information on their population status, and therefore have a significant role in setting conservation priorities (MacKenzie, 2005; Joseph et al., 2006). They are useful for a wide range of purposes from estimating changes in occurrence to identifying high conservation priority areas (Zipkin et al., 2010; Olea & Mateo-Tomas, 2011; Tilker et al. 2020). Occupancy-based diversity profiles are an important contribution to the occupancy toolkit as they allow comparing biodiversity across space and time while accounting for imperfect and varying detection. Specifically, these profiles can be used to: (1) monitor the diversity of a community over time and to evaluate the effectiveness of management / conservation efforts, and (2) compare general patterns of diversity according to different habitat, disturbance, or trait regimes, helping to set conservation priorities. Incorporating this approach into conservation should improve biodiversity assessments of species and communities.

## Acknowledgements

We would like to thank the German Federal Ministry of Education and Research (BMBF FKZ: 01LN1301A; FKZ: 01LC1703A) and the Leibniz Institute for Zoo and Wildlife Research for funding this research. SLM and MJS were funded by the UK Natural Environment Research Council (NERC; NE/K016407/1). SLM was supported by a PhD scholarship jointly funded by University of Kent and NERC. We thank the Sabah Biodiversity Council, Sabah Forest Department, Yayasan Sabah, Sime Darby, Benta Wawasan, Sabah Softwoods and Innoprise Foundation for permitting site access. We are grateful to David Coombs and Tom Swinfield for providing LiDAR data used as covariates in the occupancy model.

## Data accessibility

Scripts and model code are available on github (https://github.com/jabrams23/occudiversity). For access to the data used in this study please contact the corresponding author.

## Conflict of interest

The authors declare no conflict of interests.

## Authors’ contributions

JFA, RS, and AW conceived the idea of the study. JFA performed the simulation study analysis. SLM and MJS collected the data for the case study and generated covariate information. JFA performed the occupancy analysis for the case study. JFA lead the writing of the manuscript. All authors read, contributed to, and approved the final manuscript.

## Notes

### Competing Interest Statement

The authors have declared no competing interest.

