## Supporting Information for "Capturing biodiversity complexities while accounting for imperfect detection: the application of occupancy-based diversity profiles"

***Full code for the data simulation, model, and diversity profiles can be found on github under https://github.com/jabrams23/occudiversity.***

### Forest degradation and community simulation

We simulated a habitat covariate representing "habitat disturbance" (where 0 represents undisturbed forest and 5 represents complete deforestation) for each landscape (Fig. 1A) by drawing random samples from a multivariate normal distribution. To increase realism by simulating nonrandom habitat, we explicitly included spatial autocorrelation in the simulation of the habitat covariate by using the distance matrix as our variance-covariance matrix and a decay function with a decay constant $\phi$ to specify how the relationship to other cells changes with distance (Fig. S1). The five landscapes constitute a habitat degradation gradient representative of three different logging regimes and two "patchy" landscapes that simulate activities such as compartmental logging: (1) no disturbance, (2) patchy low disturbance (i.e. low impact logging restricted to a few logging compartments), (3) low disturbance across the entire area (i.e. low impact logging conducted throughout the logging concession), (4) patchy high disturbance (i.e. conventionally logging restricted to a few logging compartments), and (5) high disturbance across the entire area (i.e. conventional logging conducted throughout the logging concession).

We then simulated the cell-level abundance of forty virtual species distributed within our 5 landscapes. We simulated abundance to generate "true" abundance-based diversity profiles. Abundance was simulated for each of the forty species, *i*, at grid cell *j* (*j* = 1, 2,...,$n_T$) in each of our five landscapes under the following model:

$N_{ij} \sim Poisson(\lambda_{ij})$ *True abundance*

$log(\lambda_{ij}) = \beta0_i + \beta1_i * habitat_j$ *Expected abundance*

Where the species-specific intercepts ($\beta0$) and dependence on the habitat covariate ($\beta1$) are simulated as normally distributed with community hyperparameters (Fig. S2).

$\beta0_i \sim Normal(\mu_0, \sigma_0^2)$

$\beta1_i \sim Normal(\mu_1, \sigma_1^2)$

We set the average response of species to habitat disturbance ($\mu_1 = -2$) as negative since most forest-adapted species will respond to logging negatively (Fig. S2). We allowed this to vary, generating a community of species with mostly negative responses, but with few species that responded positively to habitat disturbance. Average species abundance per species per grid cell at habitat=0 (i.e. $\mu_0$) was 1.

We then generated detection/non-detection data by simulating systematic repeated sampling of the community. For each landscape, we picked 100 sampling points in a grid spaced 2 km apart. At each point, we then simulated the observation process with imperfect detection:

$y_{ijk} \sim Binomial(N_{ij}, p_{ijk})$ *Observed count data*

$logit(p_{ijk}) = \alpha 0_i$ *Detection probability*

where $y_{ijk}$ is the observed count data and $p_{ijk}$ Is the probability of detection. We repeated the observation process for 10 occasions, *k*. The average detection probability for the community was set to 0.5. Species-specific intercepts for detection, $\alpha 0$, were drawn from a normal distribution describing the community ($\mu_\alpha = 0, \sigma_\alpha = 1$).

Finally, we reduced the observed count data *y* to detection/non-detection data. We chose to represent the detection process in two steps (generating counts, then reducing these to binary detection/non-detection data) to reflect how data from typical non-invasive survey methods, such as bird point counts or camera-trapping, are prepared for occupancy modeling. We compared the performance of the occupancy-based diversity profiles to true community abundance diversity profiles. All calculations were carried out in R version 3.6.0 (R Core Team, 2019).

**Model description**

We adopted the hierarchical formulation of occupancy models by Royle & Dorazio (2008) extended to a community occupancy model (Dorazio & Royle, 2005; Dorazio et al., 2006). The first level of the model represents the true community states ($w$) of all species (*i*) in each landscape's (*s*) community (i.e., whether a species *k* is part of community *s*):

$$w_{ks} \sim Bernoulli(\Omega)$$

The second level describes the ecological process determining the occurrences (*z*) of species at sampling points (*j*), governed by the local probability of occupancy ($\psi$) and conditional on *w*:

$$z_{ij}|w_{is} \sim Bernoulli(w_{is} * \psi_{ij})$$
$$logit(\psi_{ij}) = \beta 0_{is} + \beta 1_i * habitat_j$$

The third level describes the observation process, conditional on *z*, governed by the detection probability (*p*), modeling the detection history (*y*) at each point *j* and occasion, *k*:

$$y_{ijk}|z_{ij} \sim Bernoulli(z_{ij} * p_{ijk})$$
$$logit(p_{ijk}) = \alpha 0_{is}$$

The species-specific models are linked by assuming that species-specific parameters come from a common underlying distribution, governed by community hyperparameters. Note that in the simulation study, effort is constant across occasions, but variation in *p* induced by variation in effort can be modeled by including effort as a covariate in the logit-linear predictor of *p*.

To analyze our simulated data, following common practice in analyzing field data, we modelled occupancy probability as having species-specific random intercepts, $\beta 0_{ks}$, with landscape specific (indicated by *s* indexing) hyperparameters ($\mu_{\beta 0,s}, \sigma_{\beta 0,s}$), to allow for different baseline occupancy in the reserves and among species:

$$\beta 0_{ks} \sim Normal(\mu_{\beta 0,s}, \sigma_{\beta 0,s})$$

We further modelled species-specific effects on occupancy of the simulated habitat "disturbance" covariate ($\beta 1$). Detection probability included a species-specific random intercept with landscape specific hyperparameters, to allow for differences in baseline detection among reserves. In our case, the different landscapes had different abundances of animals, which leads to differences in species-level detection (Royle & Nichols, 2003). The formal model description can be found in the supporting information. We implemented the model in a Bayesian framework using JAGS (Plummer, 2003) accessed via the R packages *rjags* (Plummer, 2019). We ran three parallel Markov chains with 250,000 iterations, of which we discarded 50,000 as burn-in, and we thinned the remaining iterations by 20 to make the output more manageable. We assessed chain convergence using the Gelman-Rubin statistic (Gelman et al., 2004). Values under 1.1 indicate convergence, and all parameters in our model had a Gelman-Rubin statistic <1.1. We tested whether the model adequately fit the data by calculating a Bayesian p-value (Gelman et al., 1996).

We used a similar model structure for the analysis of the empirical dataset. The full community occupancy model had the following parameterization:

$$w_{is} \sim Bernoulli(\Omega)$$
$$z_{ij} \sim Bernoulli\left(w_{is} * \psi_{ij}\right)$$
$$logit(\psi_{ij}) = \alpha_{i,site[j]} + \beta 1_i \mathrm{AGCD}_j + \beta 2_i \mathrm{FC}_j + \beta 3_i \mathrm{RRW}_j + \varepsilon_{block[j]}$$
$$\alpha_{i,site[j]} \sim Normal(\mu_{\alpha,site[j]}, \sigma_{\alpha,site[j]})$$
$$\beta 1_i \sim Normal(\mu_{\beta 1}, \sigma_{\beta 1})$$
$$\beta 2_i \sim Normal(\mu_{\beta 2}, \sigma_{\beta 2})$$
$$\beta 3_i \sim Normal\left(\mu_{\beta 3}, \sigma_{\beta 3}\right)$$
$$\varepsilon_{block[j]} \sim Normal(\mu_{\varepsilon}, \sigma_{\varepsilon})$$
$$y_{ijk} \sim Bernoulli\left(p_{ijk}\right)$$
$$logit(p_{ijk}) = \alpha.p_{i,reserve[j]} + \mathrm{\beta.t}_i \mathrm{time}_{jk} + \mathrm{\beta.d}_i \mathrm{date}_{jk}$$
$$\alpha.p_{i,reserve} \sim Normal(\mu.p_{\alpha.p,reserve}, \sigma.p_{\alpha.p,reserve})$$
$$\beta.t_i \sim Normal(\mu.p_{\beta.t}, \sigma.p_{\beta.t})$$
$$\beta.d_i \sim Normal(\mu.p_{\beta.d}, \sigma.p_{\beta.d})$$

In the above formulas: $z_{ij}$ is the true occupancy state (0 or 1) of species *i* at site *j*; $\psi_{ij}$ is the respective occupancy probability; $\alpha$ is the intercept of the logit-linear predictor of occupancy probability, indexed by species and reserve, $\beta 1$- $\beta 3$ are the coefficients for above ground carbon density, forest cover and riparian width, respectively. Species specific intercepts and coefficients come from a Normal distribution with community means ($\mu_{\alpha,1}$ - $\mu_{\alpha,3}$ and $\mu_{\beta 1}$ - $\mu_{\beta 3}$ for coefficients) and standard deviations ($\sigma_{\alpha,1}$ - $\sigma_{\alpha,3}$ and $\sigma_{\beta 1}$ - $\sigma_{\beta 3}$ for coefficients). Following, $y_{ijk}$ are the observations (0 or 1) of species *i* at site *j* at occasion *k*; $p_{ijk}$ are the respective detection probabilities; α.p is the intercept of the logit-linear predictor of detection probability, indexed by species and reserve; *β.t* is the effect of time of day on detection probability, and *β.d* is the effect of date on detection probability. Species specific detection intercepts and *β.d* and *β.t* come from a Normal distribution with community means ($\mu.p_{\alpha.p,1}$ – $\mu.p_{\alpha.p,3}$, and $\mu.p_{\beta.t}$ and $\mu.p_{\beta.d}$ and for *β.t* and *β.d*, respectively) and standard deviations ($\sigma.p_{\alpha.p,1}$ – $\sigma.p_{\alpha.p,3}$, and $\sigma.p_{\beta.t}$, and $\sigma.p_{\beta.d}$ for *β.t* and *β.d*, respectively).

**Rank order of site diversity**

To evaluate how well occupancy-based diversity profiles were able to order landscapes by site diversity rank, we compared them to the diversity ranking in the true abundance-based profiles. Since not all communities were different in the true abundance based profiles (e.g. no disturbance and patchy low disturbance landscapes both had an average species richness of 40 (Table 1) and could, therefore, not be distinguished in the true-abundance based profiles), we first determined how any sites we can reliably distinguish. To do so we used the results of the true abundance-based simulations and checked which landscapes could be distinguished in 95% of the 100 simulations for R, H', and D. We were able to distinguish 3, 4, and 3 of the 5 sites in the true abundance profiles for R, H', and D, respectively (Table S1).

***Table S1:*** *Percentage of the true abundance-based simulations where pairwise diversity could be distinguished for R, H', and D. We were able to distinguish between 3, 4, and 3 of the 5 sites in the true abundance profiles for R, H', and D, respectively.*

***q* = 0 (R)**

| | Site 1 | Site 2 | Site 3 | Site 4 |
|---|---|---|---|---|
| Site 2 | **0%** | - | - | - |
| Site 3 | 100% | 100% | - | - |
| Site 4 | 100% | 100% | **87%** | - |
| Site 5 | 100% | 100% | 100% | 100% |

***q* = 1 (H')**

| | Site 1 | Site 2 | Site 3 | Site 4 |
|---|---|---|---|---|
| Site 2 | 97% | - | - | - |
| Site 3 | 100% | 100% | - | - |
| Site 4 | 100% | 100% | **72%** | - |
| Site 5 | 100% | 100% | 95% | 97% |

***q* = 2 (D)**

| | Site 1 | Site 2 | Site 3 | Site 4 |
|---|---|---|---|---|
| Site 2 | 98% | - | - | - |
| Site 3 | 99% | 100% | - | - |
| Site 4 | 100% | 100% | **44%** | - |
| Site 5 | 100% | 100% | **82%** | **78%** |

**Construction of taxonomy, diet, and phylogeny similarity matrices**

Following Leinster and Cobbold (2012) we constructed the taxonomy-based similarity matrix by defining a similarity matrix **Z** by

$$Z_{ih} = \begin{cases} 0 \text{ if the } i\text{th and } h\text{th species are of different genera} \\ 0.5 \text{ if the } i\text{th and } h\text{th species are different but congeneric} \\ 1 \text{ if } i = h \end{cases}$$

The diet similarity matrix was constructed using a pairwise comparison of species dietary preferences. The similarity matrix for phylogeny was constructed using the phylogenetic distance tree for the 167 species obtained from https://birdtree.org/. The values were then linearly rescaled to a minimum of 0 and a maximum of 1.

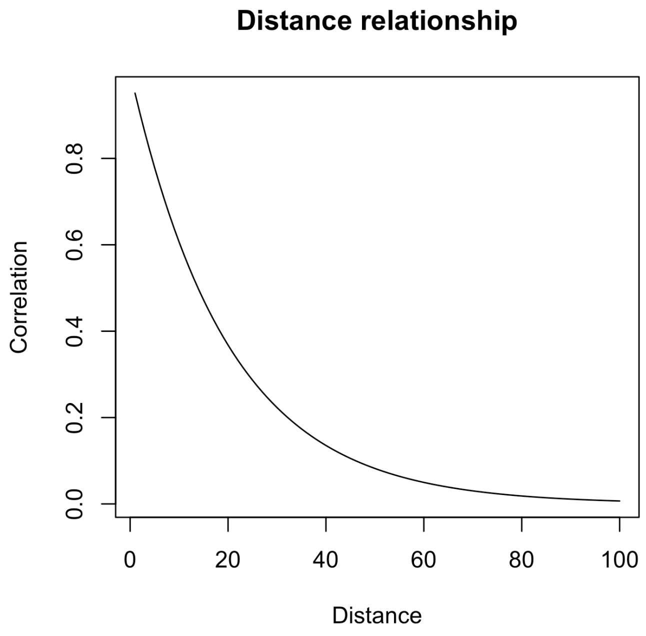


***Figure S1.*** *Distance function used to simulate virtual landscape.*

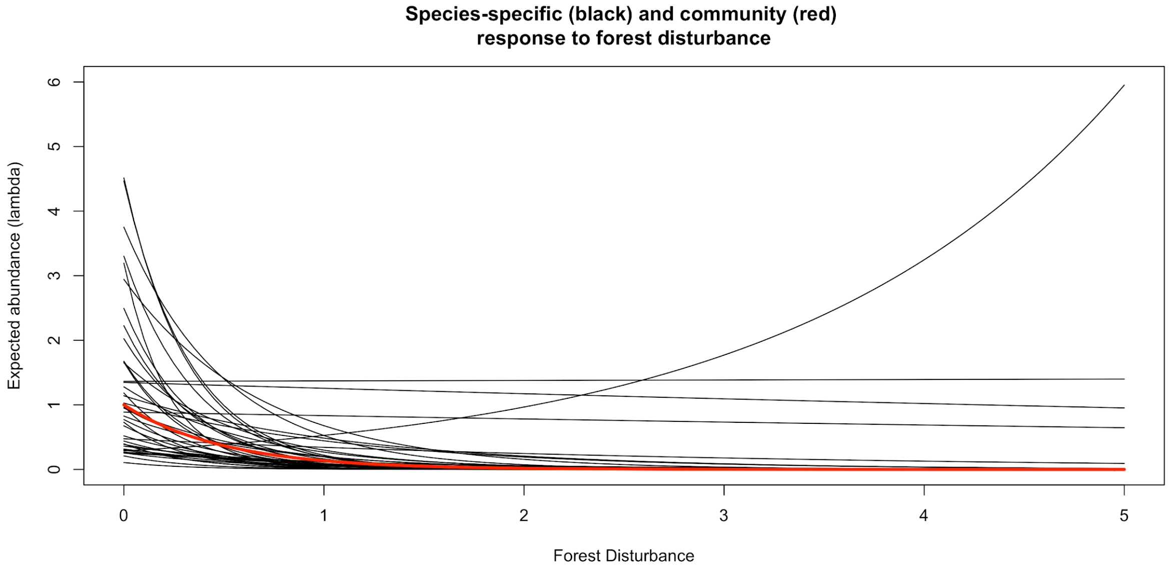


***Figure S2.*** *Simulated responses to the habitat covariate for 40 species (black lines) according to the community hyperparameter (red line).*

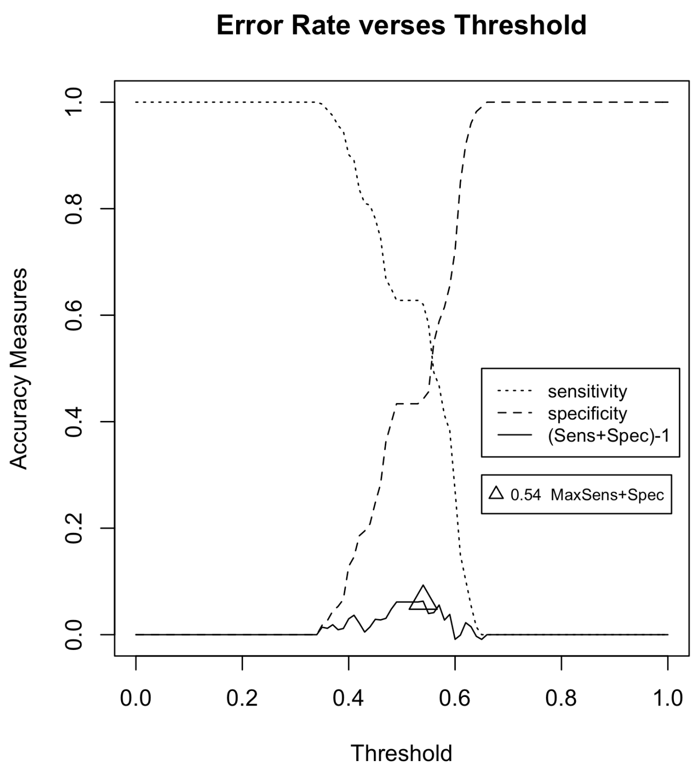


***Figure S3.*** *Example of threshold determination for one species using the* $max_{SSS}$ *method that maximizes the sum of sensitivity and specificity. This was run as a Monte Carlo simulation to produce a distribution.*

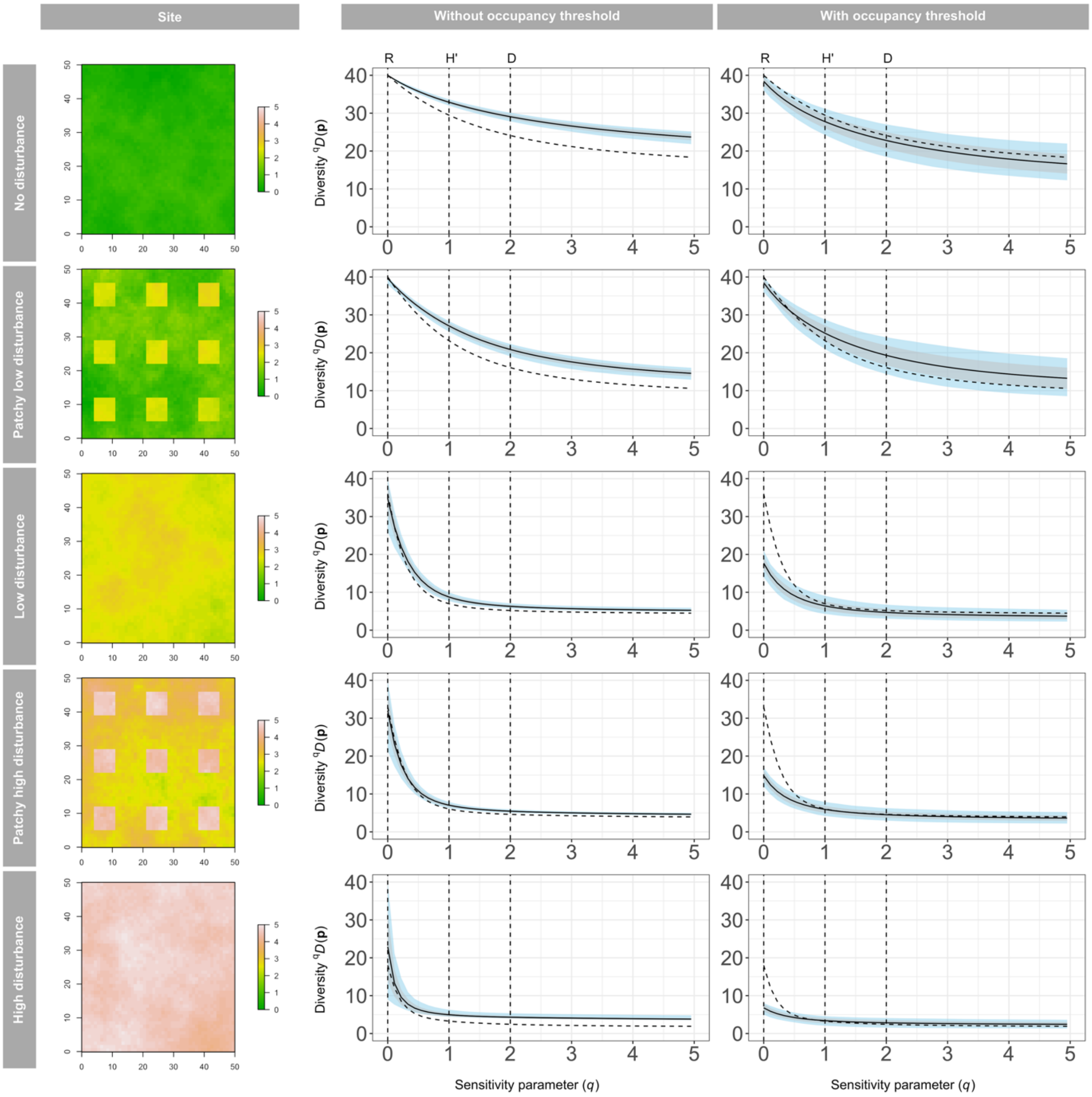


***Figure S4.*** *Results from the simulation study.* ***(A)*** *Simulated "forest quality" habitat covariate for five virtual landscapes. Diversity profiles for simulated data generated using the community occupancy predictions for the entire landscape* ***(B)*** *without thresholding and* ***(C)*** *with thresholding using the $max_{SSS}$ method for the entire study landscape (solid line, SD grey shading, 95% CI blue shading) compared to the diversity profiles generated using the true abundance across the whole landscape (dashed line).*

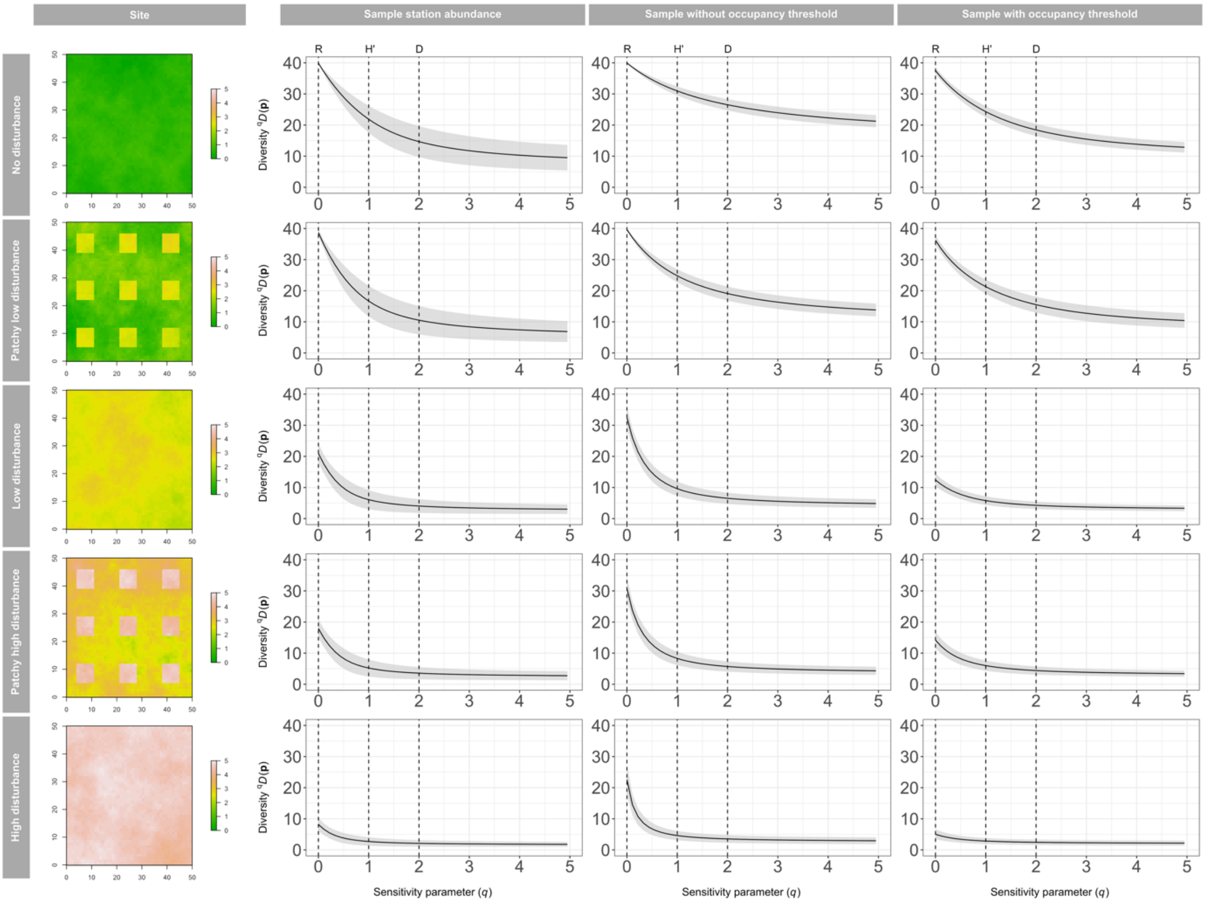


***Figure S5.*** *Results from the simulation study.* ***(A)*** *Simulated "forest quality" habitat covariate for five virtual landscapes. Average diversity profiles (solid line) for simulated data generated using data from only the 100 surveyed stations for* ***(B)*** *the simulated true abundance, and the community occupancy predictions for the entire landscape* ***(C)*** *without thresholding and* ***(D)*** *with thresholding using the* $max_{SSS}$ *method for the entire study landscape, showing the standard deviation (grey shading).*

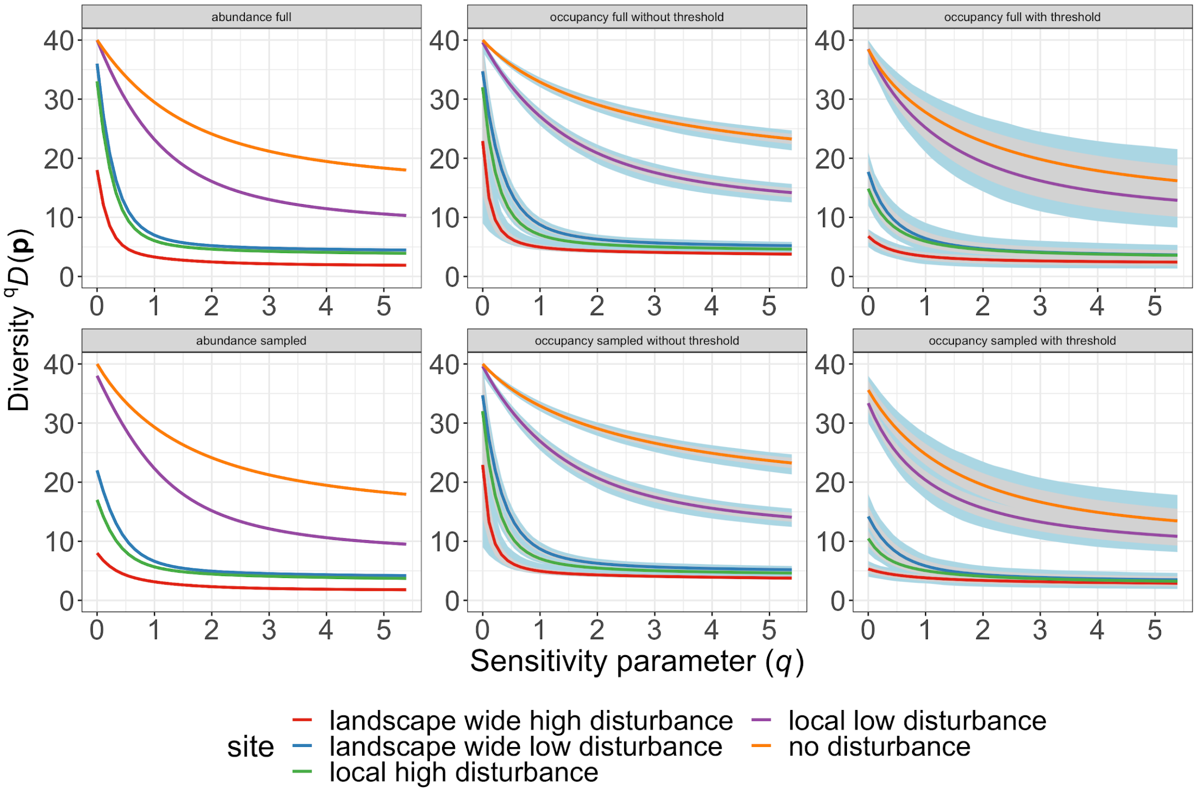


***Figure S6.*** *Comparison among landscapes of diversity profiles generated using* ***(A)*** *the true abundance across the whole landscape,* ***(B)*** *the true abundance at the 100 sample stations in each landscape,* ***(C)*** *occupancy based predictions across the whole landscape without thresholding,* ***(D)*** *occupancy based predictions at the 100 sample stations in each landscape without thresholding,* ***(E)*** *occupancy based predictions across the whole landscape with thresholding,* ***(F)*** *occupancy based predictions at the 100 sample stations in each landscape with thresholding. The occupancy-based diversity profiles are derived quantities from the Bayesian models allowing for the calculation of uncertainty*

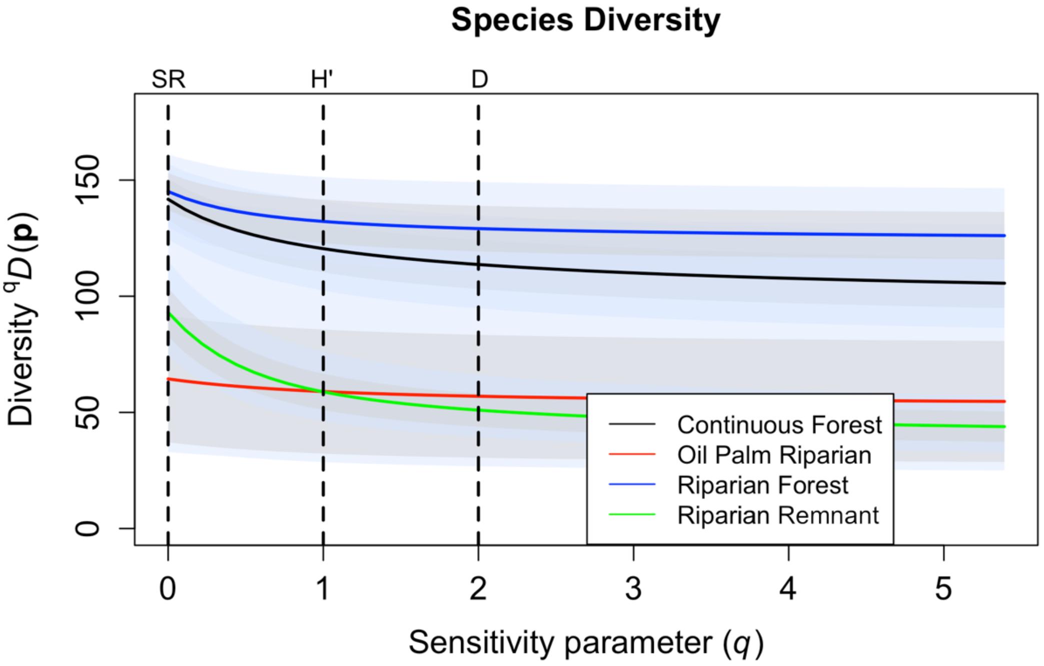


***Figure S7.*** *Diversity profiles for the empirical bird dataset with occupancy thresholding using the max$_{SSS}$ method*
